# Social interaction with an alcohol-intoxicated or cocaine-injected peer selectively alters social behaviors and drinking in adolescent male and female rats

**DOI:** 10.1101/718866

**Authors:** Danielle M. Gamble, Chloe C. Josefson, Mary K. Hennessey, Ashley M. Davis, Renee C. Waters, Brooke N. Jones, Destiny M. Belton, Nzia I. Hall, Taylor J. Costen, Cheryl L. Kirstein, Antoniette M. Maldonado-Devincci

## Abstract

**Background:** Drinking alcohol is facilitated by social interactions with peers, especially during adolescence. The importance of peer social influences during adolescence on alcohol and substance use have recently received more attention. We have shown that social interaction with an alcohol-intoxicated peer influences adolescent alcohol drinking differently in male and female rats using the demonstrator-observer paradigm. The present set of experiments analyzed the social interaction session to determine behaviors that influence alcohol drinking in adolescent male and female rats.

**Methods:** Specifically, in experiment one we determined which behaviors were altered during social interaction with an alcohol-intoxicated demonstrator and assessed changes in ethanol intake in adolescent observers. Experiment two examined changes in voluntary saccharin consumption to determine if social interaction with an alcohol-intoxicated demonstrator altered consumption of a palatable solution. In experiment three, we administered a low (5 mg/kg) or high (20 mg/kg) dose of cocaine to the demonstrator and assessed changes in the adolescent observers to determine if social interaction with a ‘drugged’ peer altered social behaviors and voluntary ethanol intake.

**Results:** We showed that social interaction with an alcohol-intoxicated demonstrator (1) decreased social play and increased social investigation and social contact in adolescent male and female observers, (2) did not alter non-social behaviors, (3) did not alter saccharin consumption and (4) increased voluntary ethanol intake in adolescent female but not male observers. When the peer was injected with cocaine (1) social play was dose-dependently decreased, (2) there were no changes in other social or non-social behaviors, and (3) voluntary ethanol intake in adolescent male and female observers was unaffected.

**Conclusions:** The present results are consistent and extend our previous work showing that social interaction with an alcohol-intoxicated peer selectively alters social behaviors and alcohol-drinking in adolescent rats. Females appear to be more sensitive to elevating effects of social interaction on voluntary ethanol consumption.

## Introduction

Underage drinking continues to be a serious health problem in the United States. According to the 2017 National Survey on Drug Use and Health, approximately 7.4 million (19.7%) individuals ages 12 to 20 reported consuming alcohol within the past month (Bose *et al.*, 2018). Moreover, 4.5 million (11.9%) of these individuals reported being binge drinkers and 932,000 (2.5%) reported being heavy drinkers (Bose *et al.*, 2018). Youth who report consuming their first drink between the ages of 9 and 11 have reported higher levels of drinking during their adolescent years (Bolland *et al.*, 2016). Similarly, individuals who report initiating alcohol use in middle school (between 6th and 8th grade), reported greater lifetime use and higher frequency of illicit substances (e.g., marijuana, cocaine, crack, amphetamines, LSD, sedatives, tranquilizers, and heroin), when compared to individuals who report initiating alcohol use in high school or later (Barry *et al.*, 2016). Together, these data indicate that adolescent alcohol use can 1) influence alcohol drinking later in life and 2) is associated with polysubstance drug use.

Beginning in adolescence, sex differences in voluntary alcohol drinking have been observed (Lancaster *et al.*, 1996), and are likely mediated by different motivations for drinking in humans. While drinking rates were overall similar among male and female teens, slightly more caster, females between 12 and 17 years reported current alcohol consumption (11% vs 8.8%) and binge drinking (6% vs 4.6%) compared to similarly-aged males (Bose *et al.*, 2018). Recent work showed that females who begin drinking at an early age drink more than any other relevant comparative age group (Bolland *et al.*, 2016). These data regarding adolescent alcohol drinking are concerning given that previous research indicates the age of initiation of alcohol use can be a significant predictor of higher alcohol and other substance abuse later in life (Irons, Iacono and Mcgue, 2015; Barry *et al.*, 2016; Bolland *et al.*, 2016). These data highlight important sex differences based on adolescent alcohol use that may influence alcohol use disorders later in life.

There have been an increasing number of studies examining the effects of social interaction, social modeling, and social influences on subsequent alcohol and substance use in humans (Beck, Thombs and Summons, 1993; Urberg *et al.*, 2003; Bot *et al.*, 2005; Mundt, 2011; Fujimoto and Valente, 2015; Deutsch *et al.*, 2017; Hallgren *et al.*, 2017; Di Guiseppi *et al.*, 2018). Outside of directly being offered a drink, indirect social modeling, such as perceived norms, is a primary way adolescents are influenced to consume more alcohol (Hallgren *et al.*, 2017; Di Guiseppi *et al.*, 2018). Having “popular” friends, high density social networks, and overestimation of peer alcohol use all influence the onset of alcohol use and increase the risk for early and increased alcohol use (Mundt, 2011; Fujimoto and Valente, 2015; Deutsch *et al.*, 2017). It has been suggested that these social environmental factors often contain peer pressure and alcohol consumption activities that encourage adolescents to drink more heavily (Rossheim *et al.*, 2017). Together, these data indicate that social factors readily influence alcohol drinking behaviors in humans.

Social influences on alcohol drinking have received attention using rodent models (Gauvin *et al.*, 1994; Juárez and De Tomasi, 1999; Hunt, Holloway and Scordalakes, 2001; Varlinskaya, Spear and Spear, 2001; Varlinskaya and Spear, 2002; Fernández-Vidal and Molina, 2004; Doremus *et al.*, 2005; Varlinskaya, Vogt and Spear, 2013; Varlinskaya, Truxell and Spear, 2015; Eade, Youngentob and Youngentob, 2016; Marcolin *et al.*, 2019). In animal models of social influence on alcohol drinking, the most commonly studied behaviors include social investigation, social play, and social contact (Varlinskaya, Spear and Spear, 2001; Varlinskaya and Spear, 2002; Maldonado-Devincci, Badanich and Kirstein, 2010; Varlinskaya, Truxell and Spear, 2015; Marcolin *et al.*, 2019). Adolescent animals seek environmental cues paired with social interactions (Calcagnetti and Schechter, 1992) and males are particularly sensitive to these effects (Maldonado-Devincci, Badanich and Kirstein, 2010). Additionally, ethanol intake is increased following social interaction more in males compared to females (Hunt, Holloway and Scordalakes, 2001; Maldonado, Finkbeiner and Kirstein, 2008). Females consume more ethanol for its socially anxiolytic effects, whereas males consume more ethanol for its social enhancing properties (Varlinskaya, Truxell and Spear, 2015). Dose-dependent changes in adolescent social behaviors have been reported, with lower ethanol doses (0.5 and 0.75 g/kg) producing social facilitation, and higher doses (2.0, 3.0, and 4.0 g/kg) inducing social inhibition (Varlinskaya, Spear and Spear, 2001). These data indicate that social interaction and alcohol produce sex-dependent changes in social behaviors and alcohol drinking in animal models.

In previous work we have shown that social interaction with an alcohol-intoxicated peer enhanced alcohol intake in adolescent male and female rats (Maldonado, Finkbeiner and Kirstein, 2008) and that only adolescent males showed a preference for environmental cues that were paired with an alcohol-intoxicated peer (Maldonado-Devincci, Badanich and Kirstein, 2010). The present experiments examined the behavioral components that may mediate sex differences following social interaction with an alcohol-intoxicated peer on changes in ethanol consumption (Experiment 1) or a saccharin-sweetened solution (Experiment 2). Experiment 3 examined whether this effect is specific to social interaction with an alcohol-intoxicated peer, or if similar effects would be observed following social interaction with a cocaine-injected peer.

## Methods and Materials

### Subjects

Eighty-two male and female (Experiment 1; 41 observers and 41 demonstrators), 78 male and female (Experiment 2; 39 observers and 39 demonstrators), and 144 male and female (Experiment 3; 72 observers and 72 demonstrators) Sprague-Dawley rats (Harlan Laboratories, Indianapolis, IN) derived from established breeding pairs at the University of South Florida, Tampa were used as subjects for the present set of experiments. Litters were sexed and culled to 10 pups per litter on postnatal day (PND) 1, with the date of birth designated as PND 0. Each litter was culled to six males and four females whenever possible. Extra pups were used in other ongoing experiments. Pups remained with their respective dams until PND 21 when pups were weaned and pair-housed with same-sex littermates, which served as the dyad pairs. Animals were maintained on a 12:12 hour light: dark cycle, with lights on from 0700 to 1900 hour, in a temperature-and humidity-controlled vivarium. Animals were allowed ad libitum access to food and water in their home cage. No more than one male or one female pup per litter was used in any given condition. Animals were used in the experiments from PND 28-30. In all respects, maintenance and treatment of the animals were within the guidelines for animal care by the National Institutes of Health.

### Apparatus

Procedures were similar to our previously published work (Maldonado, Finkbeiner and Kirstein, 2008). The non-experimental animals (demonstrators) were administered ethanol or water (Experiments 1 and 2); or cocaine or saline (Experiment 3). The experimental animals (observers) were assessed for changes in voluntary sweetened ethanol or saccharin intake. In Experiments 1 and 2, demonstrators were intragastrically administered water or 12.6% v/v ethanol mixed in water by using 12-cm lengths of polyethylene tubing (PE-50) attached to a 21.5 gauge blunted needle tip and a 5 mL disposable syringe. In experiment 3, the demonstrators were intraperitoneally (i.p.) administered 0.0, 5.0 or 20.0 mg/kg cocaine dissolved in saline (0.9% NaCl) at a dose of 1.0 ml/kg.

Assessment for voluntary intake by observers of ethanol, saccharin, and water was performed with 500 mL glass bottles with double ball-bearing tips. All bottles were weighed to the nearest 0.1 g prior to introducing and after removal of the bottles in the cage. Spillage was accounted for by placing the bottles in an empty holding cage overnight. The difference in weight was subtracted from each animal’s intake score to account for spillage that may have occurred overnight. Ethanol intake was expressed as grams of ethanol consumed per kilogram of body weight (g/kg). Saccharin intake was expressed as mL of saccharin consumed per kilogram of body weight (ml/kg).

### Procedure

All procedures were similar for experiments one and two. Procedures for experiment three were modified for the time point of cocaine administration based on previous data collected indicating maximal peak dopamine (DA) levels occurred in early adolescent rats at approximately 15 min post-injection (Badanich, Adler and Kirstein, 2006). Rats were handled for five min on PND 28 and 29 between 1730-1900.

### Experiment 1: Social interaction with an alcohol-intoxicated peer and voluntary ethanol intake

Briefly, on PND 30, at approximately 1730, each rat was weighed and socially isolated for 30 min. The demonstrator was moved to a holding cage, while the observer remained in the home cage. After 30 min of social isolation, the demonstrator was intragastrically administered 1.5 g/kg ethanol or an isovolumetric administration of water and remained socially isolated for an additional 30 min. After the 60 min of social isolation, the demonstrator and observer were reunited in the home cage and allowed to socially interact for 30 min, which was video-recorded and scored offline. After the 30 min social interaction session, the demonstrator was removed and the observer was given access to sweetened ethanol (5.6% v/v ethanol sweetened with 0.1% w/v saccharin) and water in two drinking bottles overnight for 13 hours.

### Experiment 2: Social interaction with an alcohol-intoxicated peer and voluntary saccharin intake

All procedures for experiment 2 were identical to experiment 1, except that observer rats were given two-bottle choice access between a saccharin-sweetened solution (0.1% w/v saccharin in water) and a bottle containing unsweetened water, as described above.

### Experiment 3: Social interaction with a cocaine-injected peer and voluntary ethanol intake

On PND 30, at approximately 1730, each rat was weighed and socially isolated for 45 min. The demonstrator was moved to a holding cage, while the observer remained in the home cage. After 45 min of social isolation the demonstrator was administered cocaine (5.0 or 20.0 mg/kg; i.p.) or saline (0.0 mg/kg) and remained in social isolation for an additional 15 min. After the 60 min of social isolation, the demonstrator and observer were reunited in the home cage and allowed to socially interact for 30 min, which was video-recorded and scored offline. After the social interaction session, the demonstrator was removed and the observer was given access to sweetened ethanol (5.6% v/v ethanol sweetened with 0.1% w/v saccharin) and water, as described above.

### Social Behaviors

Several social behaviors were assessed offline from the recorded videos. These included social investigation, social contact and social play, using definitions derived from Varlinskaya and colleagues (2001). Social investigation was defined as the observer sniffing the demonstrator. Social contact was defined as climbing over or under the demonstrator and social grooming. Social play included chasing, pinning, pouncing, and rough and tumble play behavior toward the demonstrator (Varlinskaya et al., 2001).

### Nonsocial behaviors

Nonsocial behaviors quantified, included self-grooming and rearing. Self-grooming was defined as the observer engaging in any grooming behavior toward the observer’s own body while not engaging in social activity with the demonstrator (Kalueff *et al.*, 2016). Rearing was defined as the observer standing on its hind legs in an attempt to explore the environment, which included sniffing the air and served as a measure for environmental exploration (Lever, Burton and O’Keefe, 2006).

### Design and Analyses

Data were analyzed separately for each experiment using a two factor between subjects design ANOVA. Experiments 1 and 2 were analyzed using a two-factor between subjects ANOVA with Peer (2; Intoxicated or Non-intoxicated) and Sex (2; Male or Female) as factors. Experiment 3 was analyzed utilizing a two-factor between subjects factor ANOVA with Dose (3; 0.0, 5.0 or 20.0) and Sex (2; Male or Female) as factors. The dependent variables for all experiments included social behaviors (i.e., social investigation, social contact, and social play), nonsocial behaviors (i.e., self-grooming and rearing), and consumption data (preference, g/kg, or ml/g). For all behaviors, only the first 10 min of the session was quantified according to the definition in (Varlinskaya, Spear and Spear, 2001). In the presence of an interaction, subsequent posthoc comparisons (Sidak’s test, simple effects) were used to isolate effects. The level for significance was set at 0.05.

## Results

### Experiment 1: Social interaction with an alcohol-intoxicated peer and voluntary ethanol intake

#### Social Behaviors

Social behaviors were quantified during the first 10 min of the session (see Varlinskaya et al., 2001). Social play was decreased by 54.99 ± 13.63% following social interaction with an alcohol-intoxicated peer in both adolescent males and females (Figure 1A, p < 0.05) as supported by a significant main effect of Peer [F (1, 37) = 15.48, p < 0.0005]. Social interaction with an alcohol-intoxicated peer increased social investigation behavior by 24.78 ± 10.82% in both adolescent male and female rats (Figure 1B) as supported by a significant main effect of Peer [F (1, 37) = 4.69, p < 0.05]. There was a strong trend for this effect to be greater in females relative to males {Sex {F (1, 37) = 3.85, p = 0.057). The interaction failed to reach statistical significance. As depicted in Figure 1C, social interaction with an alcohol-intoxicated peer increased social contact behavior by 34.44 ± 10.35% similarly in adolescent male and female observers as supported by a significant main effect of Peer [F (1, 37) = 11.61, p < 0.005]. The main effect of Sex and the interaction between Peer and Sex failed to reach statistical significance.

**Figure 1:**
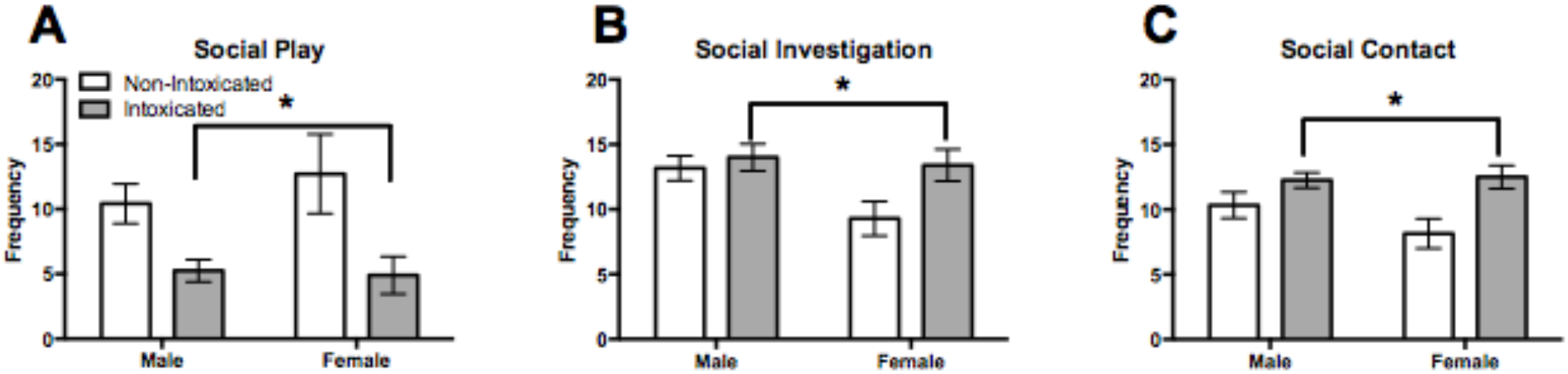
Social interaction with an alcohol-intoxicated peer differentially alters social behaviors in adolescent male and female rats. Data are presented as mean +/-SEM. As depicted in the figure, social interaction (Panel A) decreased social play, (Panel B) increased social investigation, and (Panel C) increased social contact in adolescent demonstrators that socially interacted with an alcohol-intoxicated peer compared to adolescent demonstrators that socially interaction with non-intoxicated controls. There were no sex differences in these behaviors. * Indicates a significant main effect of peer.

#### Nonsocial Behaviors

There was no change in self-grooming behavior, regardless of peer exposure or sex (Figure 2A). In general, social interaction with an alcohol-intoxicated peer increased rearing by 22.29 ± 10.20% (Main Effect of Peer [F (1, 37) = 4.59, p < 0.05] compared to those that socially interacted with a non-intoxicated peer (Figure 2B). The main effect of Sex and the Sex by Peer interaction failed to reach statistical significance.

**Figure 2:**
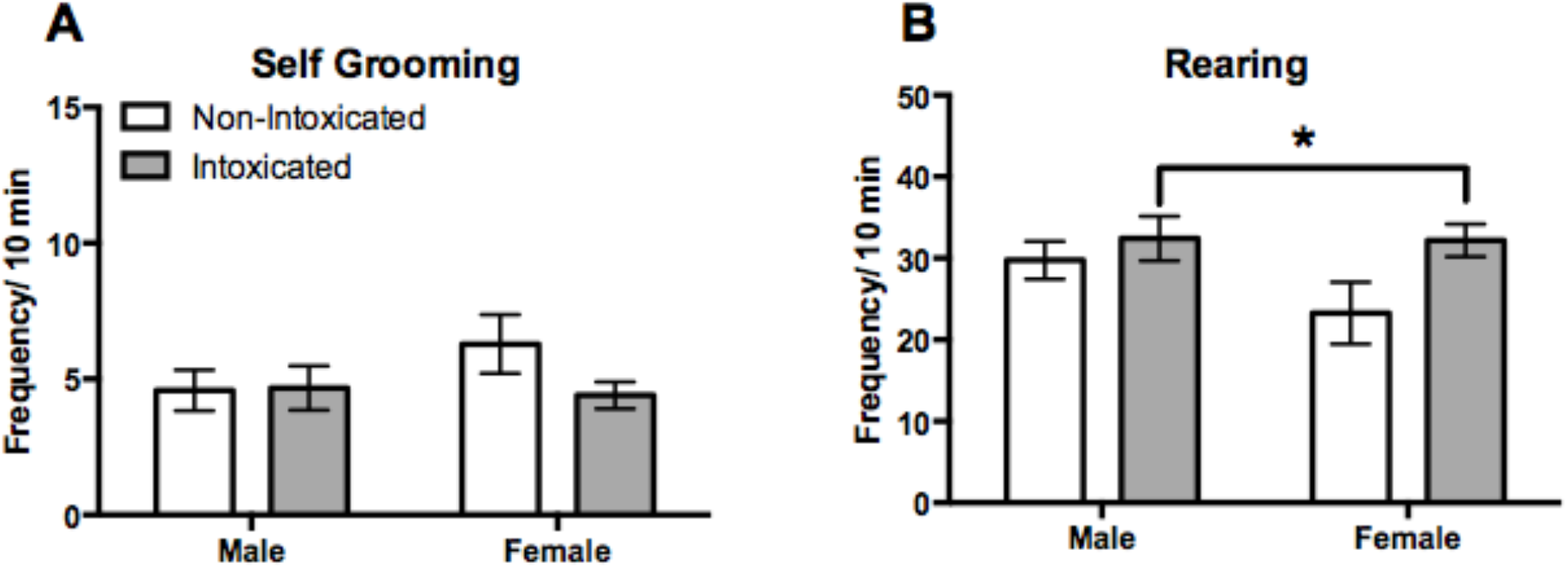
Social interaction with an alcohol-intoxicated peer differentially alters non-social behaviors in adolescent male and female rats. Data are presented as mean +/-SEM. As depicted in the figure, social interaction with an alcohol-intoxicated peer (Panel A) did not alter self-grooming, (Panel B) and increased rearing in adolescent observers compared to adolescent demonstrators that socially interacted with a non-intoxicated demonstrator peer. There were no sex differences in these behaviors. * Indicates a significant main effect of peer.

#### Ethanol Intake

Voluntary ethanol intake was assessed as grams of ethanol consumed per kilogram of body weight and as a preference ratio. Voluntary ethanol intake varied as a function of sex and peer as supported by a significant two-way interaction of Sex by Peer [F (1, 39) = 4.14, p < 0.05] and a trend for a main effect of Peer [F (1, 39) = 3.80, p = 0.058]. As shown in Figure 3A, females that socially interacted with an alcohol-intoxicated peer consumed 82.92 ± 27.70% more sweetened ethanol, when expressed as g/kg, relative to females that socially interacted with a non-intoxicated peer (p = 0.02). This effect was absent in males, regardless of peer and there were no statistically significant differences in ethanol intake (g/kg) between males and females. As depicted in Figure 3B {Sex by Peer [F (1, 39) = 5.39, p < 0.05]}, adolescent females that socially interacted with an alcohol-intoxicated peer showed higher ethanol preference relative to those that socially interacted with a non-intoxicated peer (p < 0.05), an effect that was absent in adolescent males.

**Figure 3:**
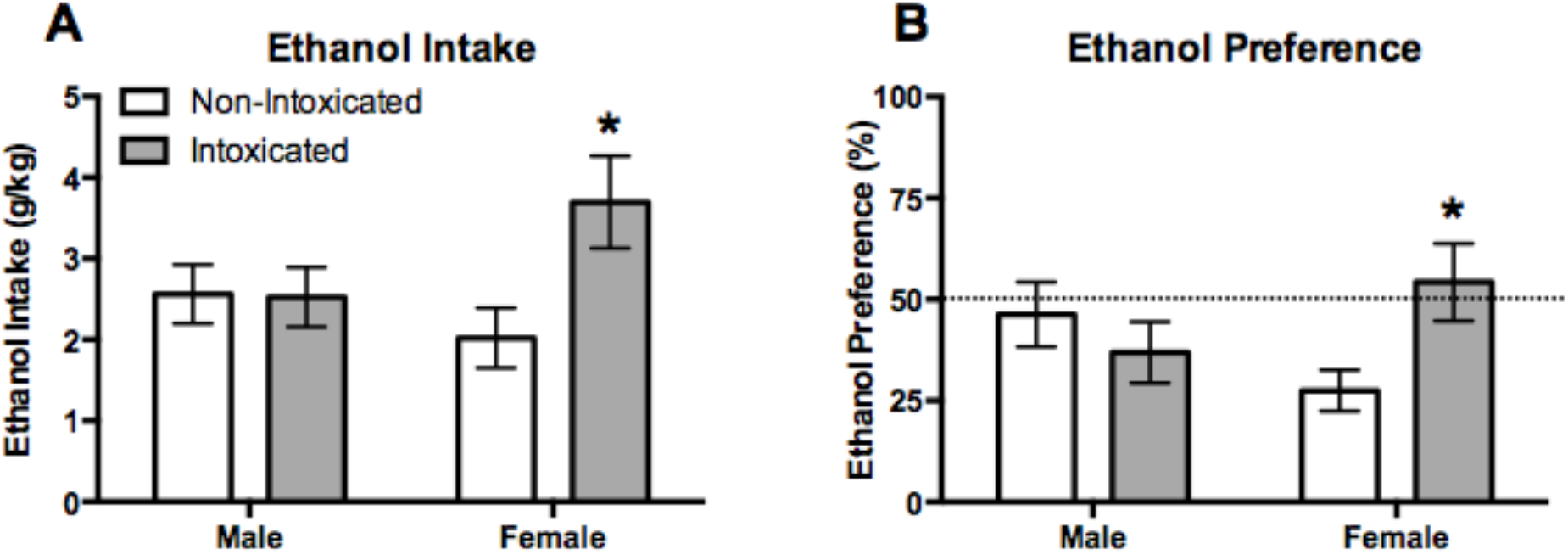
Social interaction with an alcohol-intoxicated peer increased voluntary sweetened ethanol intake (Panel A) and ethanol preference (Panel B) in adolescent female rats. Data are presented as mean +/-SEM. * Indicates social interaction with an alcohol-intoxicated peer was significantly greater than social interaction with a non-intoxicated control.

### Experiment 2: Social interaction with an alcohol-intoxicated peer and voluntary saccharin consumption

#### Social Behaviors

Similar to Experiment 1, social interaction with an alcohol-intoxicated peer decreased social play by −47.82 ± 18.3% in both males and females (Figure 4A) as supported by a Main Effect of Peer [F (1, 39) = 7.47, p < 0.01]. The main effect of Sex and Sex by Peer interaction failed to reach statistical significance. Additionally, social interaction with an alcohol-intoxicated peer increased social investigation by 25.72 ± 6.92% in both males and females (Figure 4B) as supported by a Main Effect of Peer [F (1, 39) = 13.48, p < 0.001]. The Main Effects of Sex and Peer by Sex interactions failed to reach statistical significance. Finally, social interaction with an alcohol-intoxicated peer did not alter social contact, regardless of sex (Figure 4C).

**Figure 4:**
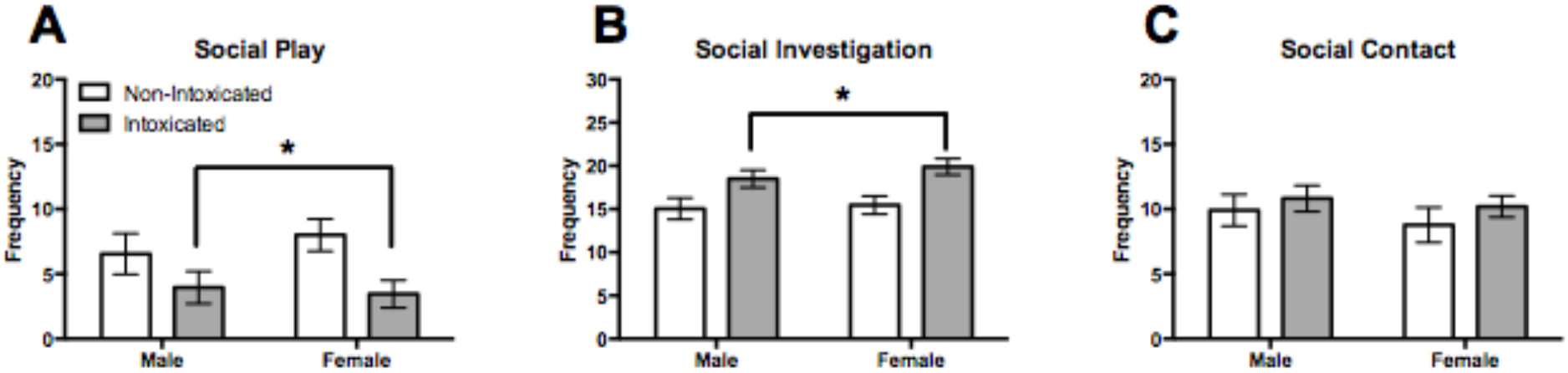
Social interaction with an alcohol-intoxicated peer differentially alters social behaviors in adolescent male and female rats. Data are presented as mean +/-SEM. As depicted in the figure, social interaction (Panel A) decreased social play, (Panel B) increased social investigation, and (Panel C) did not alter social contact in adolescent demonstrators that socially interacted with an alcohol-intoxicated peer compared to adolescent demonstrators that socially interaction with non-intoxicated controls. There were no sex differences in these behaviors. * Indicates a significant main effect of peer.

#### Nonsocial Behaviors

Self-grooming and rearing were not altered in adolescent male and female rats that socially interacted with an alcohol-intoxicated peer (Figure 5).

**Figure 5:**
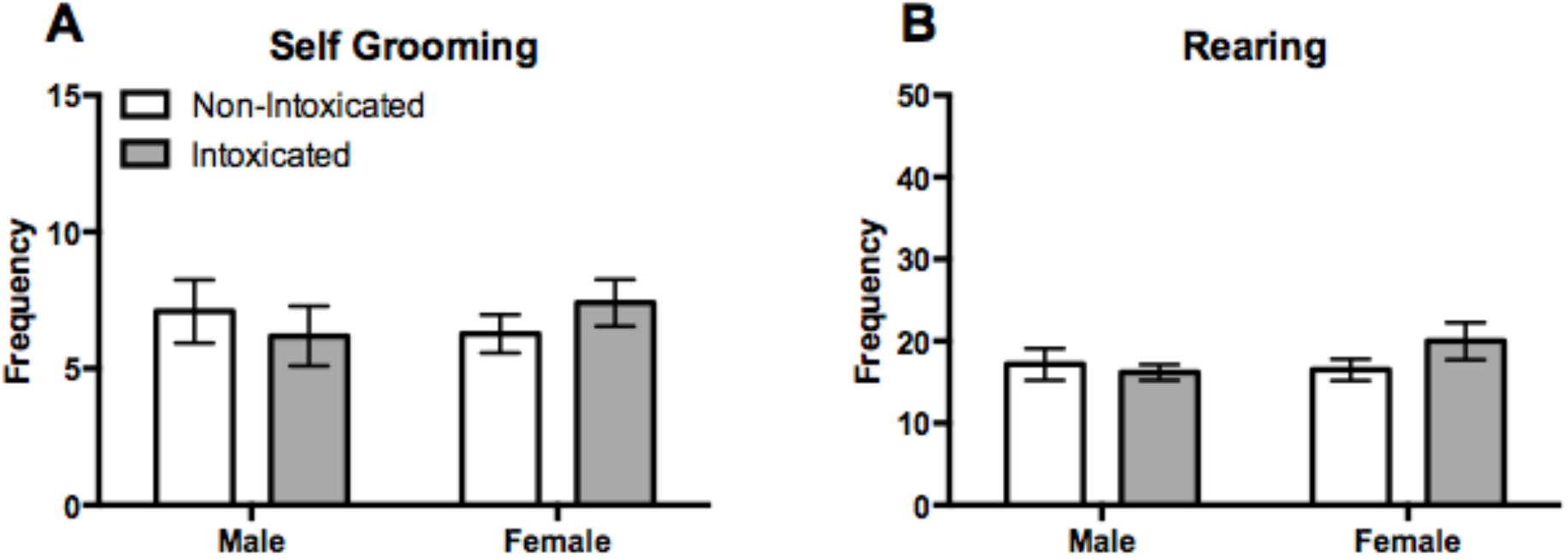
Social interaction with an alcohol-intoxicated peer did not alter the non-social behaviors of self-grooming (Panel A) or rearing (Panel B) in adolescent male and female rats. Data are presented as mean +/-SEM.

#### Saccharin Intake

Voluntary saccharin intake was assessed as milliliters of saccharin consumed per kilogram of body weight and as a preference ratio. As shown in Figure 6, there was no change in voluntary saccharin intake (Figure 6A) and no difference in preference for the saccharin solution (Figure 6B), when the observer socially interacted with an alcohol-intoxicated demonstrator.

**Figure 6:**
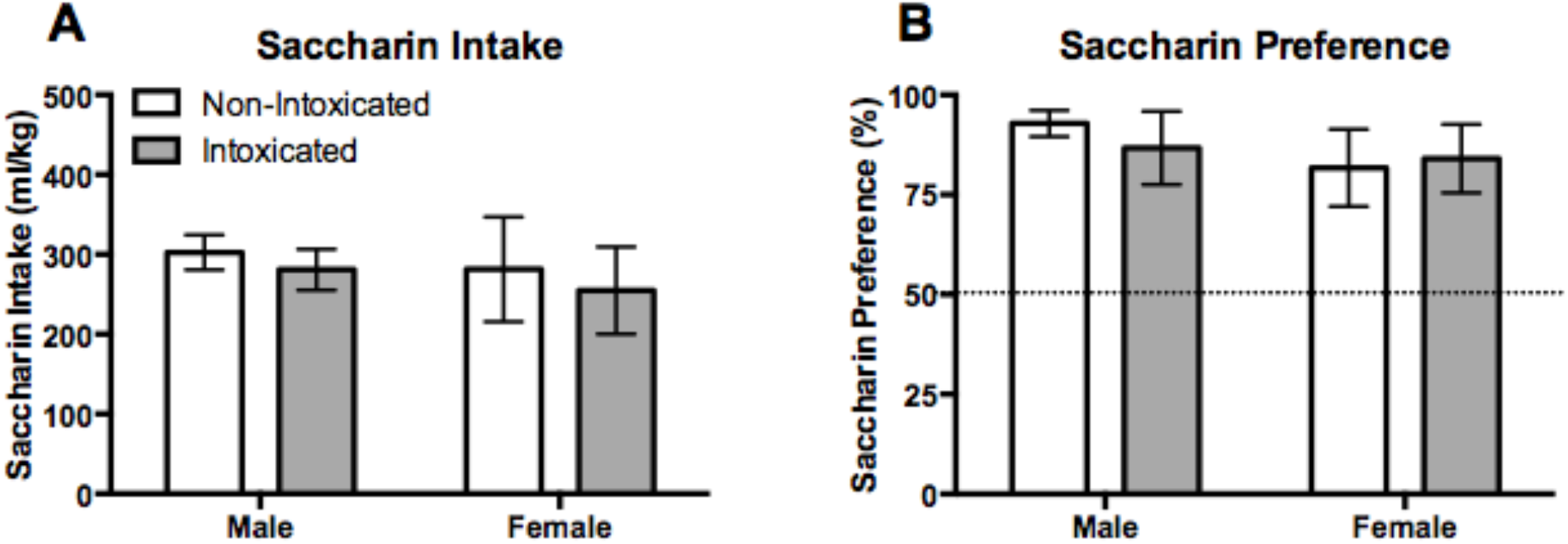
Social interaction with an alcohol-intoxicated peer did not alter the voluntary saccharin intake (Panel A) or saccharin preference (Panel B) in adolescent male and female rats. Data are presented as mean +/-SEM.

### Experiment 3: Social interaction with a cocaine-injected peer and voluntary ethanol intake

#### Social Behaviors

Social play was dose-dependently decreased in observers that socially interacted with a cocaine-injected demonstrator (Figure 7A) as supported by a main effect of Peer [F (2, 64) = 14.18, p < 0.0001]. When collapsed across Sex, social interaction was decreased by −39.41 ± 12.86% (p < 0.01) when the adolescent observer socially interacted with a peer that was administered 5 mg/kg compared to controls. Similarly, social interaction was decreased by - 69.04 ± 12.73% (p < 0.0001) when the adolescent observer socially interacted with a peer that was administered 20 mg/kg cocaine compared to controls. Additionally, social play behavior lowered by 29.63 ± 12.72% following social interaction with a peer administered 20 mg/kg cocaine compared to social interaction with a peer that was administered 5 mg/kg cocaine (p < 0.01). The main effect of Sex and the Peer by Sex interaction failed to reach statistical significance.

**Figure 7:**
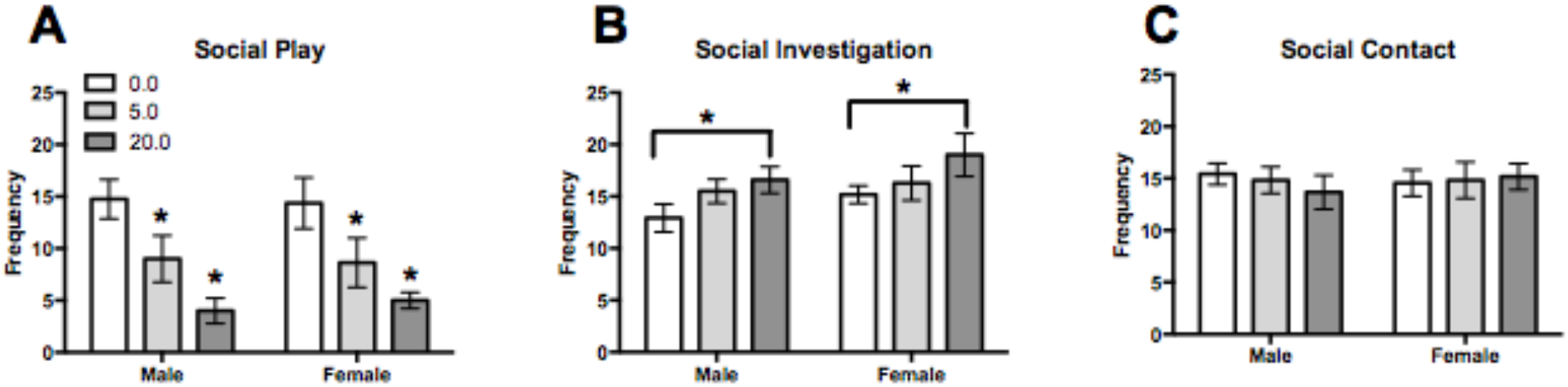
Social interaction with a cocaine-injected peer differentially alters social behaviors in adolescent male and female rats. Data are presented as mean +/-SEM. As depicted in the figure, social interaction with a cocaine-injected peer (Panel A) dose-dependently decreased social play, (Panel B) increased social investigation, and (Panel C) increased social contact in adolescent demonstrators that socially interacted with a cocaine-injected peer compared to adolescent demonstrators that socially interacted with vehicle-administered controls. There were no sex differences in these behaviors. * Indicates a significant difference from 0 mg/kg.

In general, social interaction with a cocaine-injected peer increased social investigation, as supported by a main effect of Peer [F (2, 64) = 3.33, p < 0.05]. However, posthoc analyses failed to reveal any statistically significant differences (Figure 7B). The main effect of Sex and the Peer by Sex interaction failed to reach statistical significance. Social interaction with a cocaine-injected peer did not alter social interaction in the observer (Figure 7C). The main effect of Sex, the main effect of Peer, and the Peer by Sex interaction failed to reach statistical significance.

#### Nonsocial Behaviors

In general, social interaction with a cocaine-injected peer administered 5 or 20 mg/kg cocaine increased self-grooming by 45.10 ± 17.21% or 52.61 ± 16.60%, respectively, compared to social interaction with a peer administered vehicle (0 mg/kg) (Figure 8A). Social interaction with a peer demonstrator that was administer 20 mg/kg cocaine showed 35.88 ± 14.42% higher rearing compared to social interaction with a peer administered vehicle (0 mg/kg), regardless of Sex (Figure 8B).

**Figure 8:**
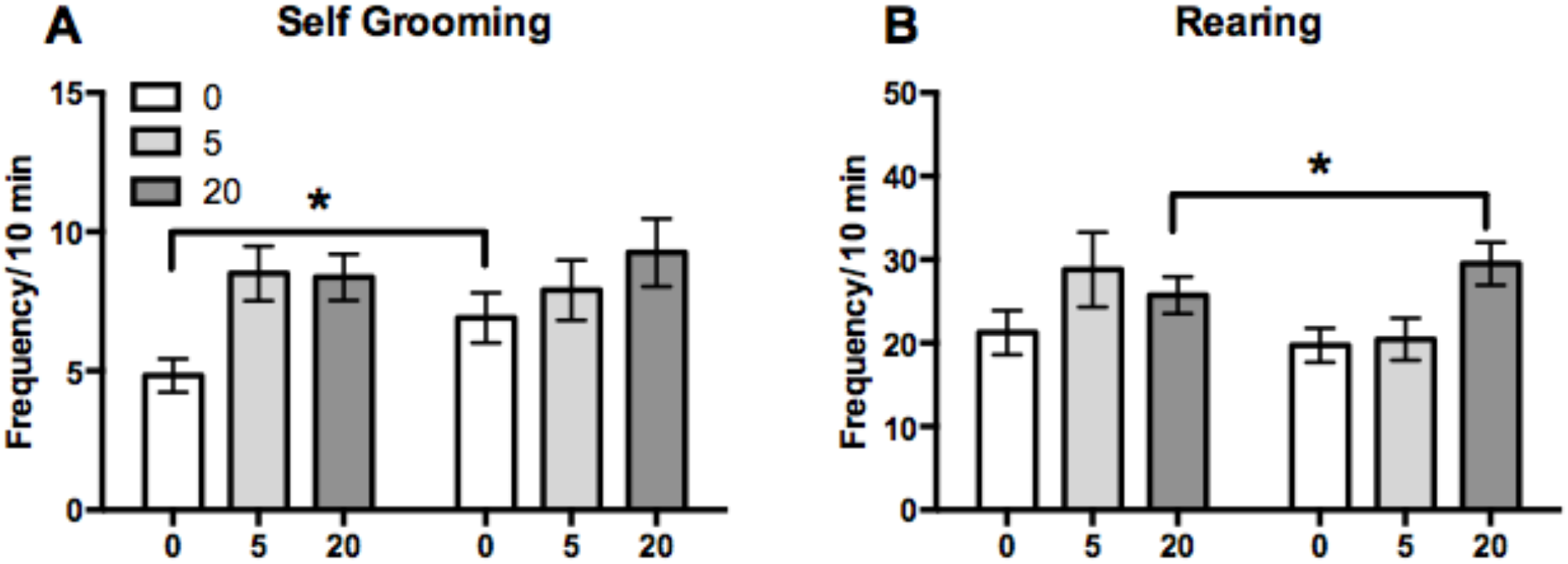
Social interaction with a cocaine-injected peer differentially increased the non-social behaviors of (Panel A) self-grooming and (Panel B) rearing. Data are presented as mean +/-SEM. There were no sex differences in these behaviors. * Indicates 0.0 is significantly different from other doses.

#### Ethanol Intake

Voluntary ethanol intake was not altered following social interaction with a cocaine-injected peer. However, females consumed 24.07 ± 10.81% more ethanol compared to males (Figure 9A) as supported by a main effect of Sex [F (1, 66) = 9.59, p < 0.005]. The main effect of Peer and the Sex by Peer interaction failed to reach statistical significance. However, there were no differences in preference for the ethanol solution following social interaction with a cocaine-injected peer, regardless of sex (Figure 9B).

**Figure 9:**
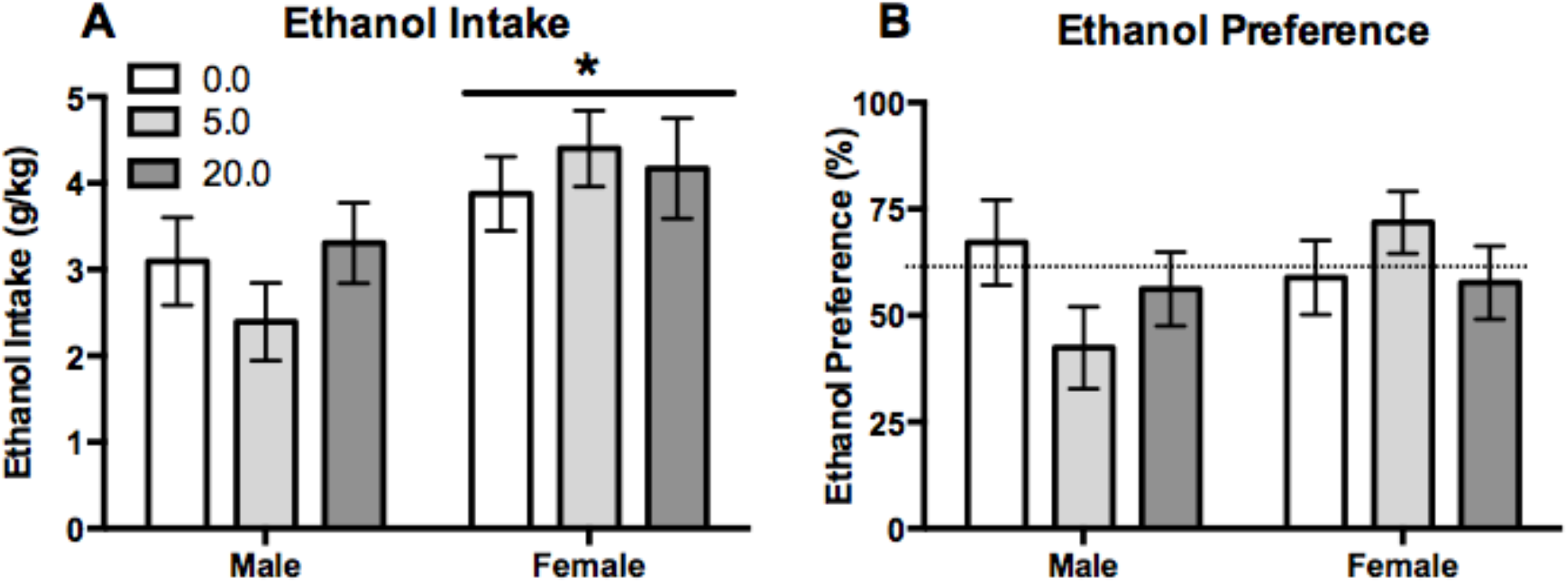
Social interaction with a cocaine-injected peer did not alter the voluntary ethanol intake (Panel A) or ethanol preference (Panel B) in adolescent male and female rats. Data are presented as mean +/-SEM. * indicates female is significantly greater than male.

## Discussion

The present work determined which aspects of the social interaction experience influenced changes in ethanol consumption following interaction with an alcohol-intoxicated or cocaine-injected peer. Results from the present set of experiments support the hypothesis that social interaction with an alcohol-intoxicated peer influenced specific social behaviors, in turn increasing voluntary ethanol intake in adolescent female rats. Previous research shows that social interaction with a peer intoxicated with alcohol alters social behavior in adolescent rats, with low doses of ethanol increased social behaviors, such as social play and social investigation, whereas high doses of ethanol decreased social behaviors (Varlinskaya, Spear and Spear, 2001; Varlinskaya and Spear, 2009). In addition, social interaction with a drinking or an intoxicated peer can alter drinking behaviors by increasing alcohol consumption in adolescent rats (Maldonado-Devincci, Badanich and Kirstein, 2010; Hallgren *et al.*, 2017). The present set of experiments conducted in adolescent rats replicated recent work in humans showing that being surrounded by a regular drinker increased the likelihood of the adolescent to drink similarly in both age-matched regular drinking adolescent siblings and age-matched regular drinking friends in both males and females (Scholte *et al.*, 2008; Hallgren *et al.*, 2017; Temmen and Crockett, 2018). The present research highlights that females are disproportionately affected by exposure to alcohol during adolescence.

### Social Behaviors

Experiment 1 was designed to replicate and extend our previous study to 1) examine which behaviors during the social interaction session were altered and 2) how this influenced subsequent ethanol consumption. In this experiment, we showed that social interaction with an alcohol-intoxicated peer increased social investigation and social contact, while simultaneously decreasing social play in both males and females. Experiment 2 was designed to manipulate the solution that was consumed following interaction with an alcohol-intoxicated peer. Similar to experiment 1, we saw increased social investigation, decreased social play, and no change in social contact. These effects were primarily driven by changes in social behaviors in the females as shown in experiments 1 and 2.

Social interaction is an important aspect of the demonstrator-observer paradigm since adolescent animals engage in distinct forms of social interaction, including play behaviors and social investigation (Varlinskaya, Spear and Spear, 2001). Previous research found that ethanol amplifies the presence of social interaction through increased play fighting in male rats administered ethanol, i.e., the demonstrator in our present set of experiments (Varlinskaya, Truxell and Spear, 2015). However, in female rats social interaction breeds social anxiety-like behaviors, potentially by creating a need for ethanol’s anxiolytic effect, which is manifested with higher ethanol intake (Varlinskaya, Truxell and Spear, 2015). The present set of work examined changes in passive social influences in behaviors of the peer that was not administered ethanol, i.e., the observer in the present set of experiments, where we observed decreased play in the observer adolescent rat in both sexes. The differences in social behaviors observed in the present set of experiments may be explained by these motivational differences and passive influences (Varlinskaya, Truxell and Spear, 2015).

Experiment 3 was designed to manipulate the type of peer the adolescent socially interacted with. In this experiment, adolescent males and females socially interacted with a peer that was administered a low or high dose of cocaine and subsequent ethanol consumption was measured. In this experiment, social interaction with a cocaine-injected peer dose-dependently decreased social play, increased social investigation, but did not change social contact behavior. These findings are consistent with the other two experiments in this study and with previous literature that found cocaine decreased social play in adolescent rats (Achterberg *et al.*, 2014) and decreased overall social interaction in adult rats (Šlamberová *et al.*, 2015). In addition to cocaine, other drugs such as amphetamines and cannabis also decreased social play in adolescent rats, i.e., the demonstrator in the present set of experiments (Trezza and Vanderschuren, 2008; Achterberg *et al.*, 2014). There were no clear sex differences driving the changes in social behaviors following interaction with a cocaine-injected peer in the present study. Together, these data indicate that social interaction with an ethanol-intoxicated or cocaine-injected peer similarly altered social behaviors, by increasing social investigation and decreasing social play.

### Non-Social Behaviors

The non-social behaviors we quantified were self-grooming and rearing during the first 10 min of the trial. This is based on previous work (Varlinskaya, Spear and Spear, 2001), where they indicated that the first 10 min of the trial was sufficient to characterize the entire social interaction trial. It is important to note that previous studies have also identified rearing as a proactive emotional coping behavior, specifically explorative escape behavior (Paul *et al.*, 2011). In Experiments 1 and 2, there were no changes in self-grooming. However, in Experiment 3, social interaction of the observer with a cocaine-injected demonstrator increased self-grooming, regardless of sex.

In Experiment 1, social interaction with an alcohol-intoxicated peer increased rearing, but this effect was absent in Experiment 2. In Experiment 3, social interaction with a demonstrator peer administered 20 mg/kg cocaine increased rearing compared to controls, similar to recent work in adult rats (Šlamberová *et al.*, 2015). Other studies have examined these and other nonsocial behaviors in relation to substance use and found that initial exposure to nicotine increased ambulation, but decreased rearing in adolescent rats (Schochet, Kelley and Landry, 2004). Together, these findings indicate that nonsocial behaviors are also affected by exposure to drug-intoxicated peers, particularly following interaction with a cocaine-injected peer.

### Ethanol Consumption

Adolescent females that socially interacted with an alcohol-intoxicated peer showed increased voluntary ethanol intake, while males showed no change in ethanol consumption. There was no change in saccharin consumption following social interaction with an alcohol-intoxicated peer, indicating that the change in ethanol consumption was not merely due to the sweetener used in the ethanol solution. This finding supports previous research that found that adolescent rats consumed elevated amounts of ethanol, independent of the saccharin sweetener (Doremus *et al.*, 2005). In addition, ethanol consumption was not altered in males or females following social interaction with a cocaine-injected peer. These data indicate that increases in ethanol consumption are specific to social interaction with an alcohol-intoxicated peer, and this relationship is not generalized to social interaction with a peer in a different state of intoxication, i.e., social interaction with a cocaine-injected peer. Previous research conducted in rats found that when both animals, the demonstrator and observer (combined exposure) were intoxicated, there was an increase in alcohol consumption, suggesting that when both peers of the dyad are intoxicated this potentiates the risk for increased adolescent alcohol consumption (Eade, Youngentob and Youngentob, 2016). Recently, it was shown that social instability stress (isolation for one hour then placed with a new cage mate daily) during adolescence (PND 30-45) increased intake of ethanol, but not sucrose, when male rats were tested during adolescence (Marcolin *et al.*, 2019), suggesting that changes in social interaction mediate overall changes in ethanol intake in adolescent males. However, these results were not supported in the current study, as changes in ethanol intake were only observed in female adolescent rats.

This current set of experiments support previous research that indicates females appear to be more sensitive to peer norms in influencing alcohol drinking (Yeh, Chiang and Huang, 2006; Dick *et al.*, 2007). Most of the changes in the social behaviors were driven by larger changes in the females that socially interacted with an alcohol-intoxicated peer compared to males that socially interacted with an alcohol-intoxicated peer. Females tend to use alcohol more to manage emotional distress and in response to peer pressure, whereas males tend to use alcohol more with social bonding and to facilitate social interaction (Thombs, Beck and Mahoney, 1993; Opland, Winters and Stinchfield, 1995; Simons-Morton *et al.*, 2001).

Much like alcohol, intake of other drugs is mediated by gender, relationships, and socialization. With cigarette smoking among adolescents, a non-smoker’s behavior may be more likely to decrease smoking behavior in a smoker (Lakon *et al.*, 2015). However, other work suggests that adolescents are more likely to use cigarettes, marijuana, or heroin if their friends are regular users, if they believe their peers are using, or as a gateway to social status and popularity (Tucker *et al.*, 2014; Filippidis, Agaku and Vardavas, 2015; Schaefer *et al.*, 2015).

Overall, the present results suggest adolescent males and females are sensitive to environmental conditions when socially interacting with an alcohol intoxicated peer. Alcohol’s ability to induce social facilitation has been one of the most important reasons reported for initiation of alcohol use in human adolescents (Beck, Thombs and Summons, 1993; Temmen and Crockett, 2018). Additionally, selection and socialization of substance using peers also has a dramatic influence on alcohol use (Borsari and Carey, 2006; Hallgren *et al.*, 2017; Temmen and Crockett, 2018). In humans, the social facilitating impact of peers seems to have a greater influence on drinking behaviors in males than females (Miller and Prentice, 1996; Borsari and Carey, 2006; Martino *et al.*, 2006). However, recent work in rodents indicates that social interaction breeds social anxiety-like behaviors in females (Varlinskaya, Truxell and Spear, 2015). Therefore, these factors may be some of the underlying conditions for the observed sex differences and social influence factors on ethanol consumption. Attempts to understand the underlying conditions in the initiation of alcohol use during adolescence may help to prevent the possibility of prolonged use into adulthood.

## Acknowledgements

This work was funded by the Department of Psychology at North Carolina A&T State University (AMD), the University of South Florida College of Arts and Sciences, USF Neuroscience Health Program, University of South Florida Department of Psychology and the Honors College.

